# Identification of the bacterial biosynthetic gene clusters of the oral microbiome illuminates the unexplored social language of bacteria during health and disease

**DOI:** 10.1101/431510

**Authors:** Gajender Aleti, Jonathon L. Baker, Xiaoyu Tang, Ruth Alvarez, Márcia Dinis, Nini C. Tran, Alexey V. Melnik, Cuncong Zhong, Madeleine Ernst, Pieter C. Dorrestein, Anna Edlund

## Abstract

Small molecules are the primary communication media of the microbial world. Recent bioinformatics studies, exploring the biosynthetic gene clusters (BGCs) which produce many small molecules, have highlighted the incredible biochemical potential of the signaling molecules encoded by the human microbiome. Thus far, most research efforts have focused on understanding the social language of the gut microbiome, leaving crucial signaling molecules produced by oral bacteria, and their connection to health versus disease, in need of investigation. In this study, a total of 4,915 BGCs were identified across 461 genomes representing a broad taxonomic diversity of oral bacteria. Sequence similarity networking provided a putative product class for over 100 unclassified novel BGCs. The newly identified BGCs were cross-referenced against 254 metagenomes and metatranscriptomes derived from individuals with either good oral health, dental caries, or periodontitis. This analysis revealed 2,473 BGCs, which were differentially represented across the oral microbiomes associated with health versus disease. Co-abundance network analysis identified numerous inverse correlations between BGCs and specific oral taxa. These correlations were present in health, but greatly reduced in dental caries, which may suggest a defect in colonization resistance. Finally, corroborating mass spectrometry identified several compounds with homology to products of the predicted BGC classes. Together, these findings greatly expand the number of known biosynthetic pathways present in the oral microbiome and provide an atlas for experimental characterization of these abundant, yet poorly understood, molecules and socio-chemical relationships, which impact the development of caries and periodontitis, two of the world’s most common chronic diseases.

**IMPORTANCE:** The healthy oral microbiome is symbiotic with the human host, importantly providing colonization resistance against potential pathogens. Dental caries and periodontitis are two of the world’s most common and costly chronic infectious diseases, and are caused by a localized dysbiosis of the oral microbiome. Bacterially produced small molecules, often encoded by BGCs, are the primary communication media of bacterial communities, and play a crucial, yet largely unknown, role in the transition from health to dysbiosis. This study provides a comprehensive mapping of the BGC repertoire of the human oral microbiome and identifies major differences in health compared to disease. Furthermore, BGC representation and expression is linked to the abundance of particular oral bacterial taxa in health versus dental caries and periodontitis. Overall, this study provides a significant insight into the chemical communication network of the healthy oral microbiome, and how it devolves in the case of two prominent diseases.

## INTRODUCTION

The human body is inhabited by rich and diverse bacterial communities, which are intimately linked to the health of the human host (1). Small molecules, which are often encoded by biosynthetic gene clusters (BGCs), are the primary means of communication in this microbial world. Recent studies suggest that the human microbiota has the potential to synthesize a myriad of exquisite small molecules, and that these small molecules serve as mediators in a variety of microbe-microbe and host-microbe interactions (2–4). These include: antibacterial activity (5), bacterial signaling (6), immune modulation (7), biofilm formation (8, 9), host colonization (10), nutrient-scavenging (11) and stress protection (12). Disruption of the finely-tuned equilibrium of the bacterial ecosystems in the human microbiome, referred to as dysbiosis, is associated with a plethora of diseases. While the mechanistic underpinnings of a shift to a dysbiotic community remain poorly understood, there is little doubt that signaling via the small molecules produced by microbial BGCs plays a critical role in the transition to dysbiosis, and associated pathogenesis (13, 14).

The human oral cavity contains an assortment of ecological niches, and as such, harbors one of the most diverse microbial populations in the human body (1, 15). Dental caries and periodontitis are two of the most common and costly chronic conditions afflicting humans, and are the result of localized dysbiosis in the oral cavity (16–20). Unlike the rest of the human digestive tract, the oral cavity is consistently exposed to the exterior environment. Therefore, an indispensable portion of the first line of defense against invading pathogens is the colonization resistance provided by a healthy oral microbiome. Indeed, dysbiosis of the oral microbiome is not only directly linked to oral diseases, but is also implicated in system-wide health (21), stressing the urgent need to unravel the underlying factors that shape and maintain a healthy human oral microbiome.

Elucidating the transmissions relayed by oral bacterial small molecules could lead to a deeper understanding of key ecological factors that set the stage for oral community succession, in health and pathogenesis. A large and growing body of literature suggests that the microbial composition and metabolic potential of the saliva and dental plaque varies significantly in healthy versus disease states (22–28). Therefore, we hypothesize that the abundance and expression of BGCs, which produce small molecules, may drive crucial bacterial interactions which contribute to health or disease. To explore this further, the biosynthetic capacity of 461 well-annotated oral bacterial genomes was investigated, and an enormous diversity of BGCs was revealed. In addition, sequence reads from 294 publicly available metagenomes and metatranscriptomes, which were associated with health, dental caries, or periodontitis, were mapped to these novel oral BGCs. This analysis identified 2,473 biosynthetic pathways which were differentially represented in health versus disease. In addition, the BGC content in salivary metagenomes obtained from 24 healthy children and 23 children with dental caries was analyzed. A Bayesian network approach was employed to identify both positive and inverse correlations between BGCs and bacterial taxa, which revealed differentially abundant signaling networks and species in health compared to dental caries. Overall, this study provides a significant insight into the chemical communication network of the healthy oral microbiome, and how it devolves in the case of dental caries and periodontitis.

## RESULTS AND DISCUSSION

### The human oral microbiome encodes thousands of diverse BGCs from an array of species

To explore the metabolic capacity of the human oral microbiome in-depth, a comprehensive pipeline for mining bacterial genomes was established, utilizing antiSMASH infrastructure v4 (accessible at https://antismash.secondarymetabolites.org/) (29), including MultiGeneBlast (30). An oral bacterial genome sequence database was assembled to include a total of 461 well-curated and annotated bacterial genomes, representing 113 unique bacterial genera and 298 taxonomically unique species, as well as 72 taxa unclassified at the species level (Table S1). Genomes were selected based on their completeness and level of annotation. A single genome sequence for each bacterial species was included to circumvent the overrepresentation of BGCs from bacteria with a high number of genome representatives. Indeed, in a previous bioinformatics study of 169 *S. mutans* genomes, ~1,000 putative BGCs were identified, revealing an incredible potential to produce small molecules within one bacterial species (31). Therefore, it should be noted that the estimated BGC diversity reported here is likely underestimated. Clearly, strain-level diversity is important to explore in future studies. However, this will require extensive genome sequencing, since to-date most oral bacterial species lack multiple reference genomes. By applying the genome-mining pipeline described above, a total of 4,915 BGCs of known and unknown types were identified (Table S1). BGCs annotated as fatty acid synthases, which are often involved in primary metabolism, were excluded. Approximately 50% of the identified BGCs were of an unknown class, congruent with the observations of other efforts to identify BGCs (Table S1)(2). The remaining 50% of BGCs (2,250) shared sequence similarities with an extensive range of previously characterized BGC classes, which is likely reflective of the high taxonomic diversity observed within the oral cavity as compared to many other body sites (1) (Fig. 1A).

**FIG 1:**
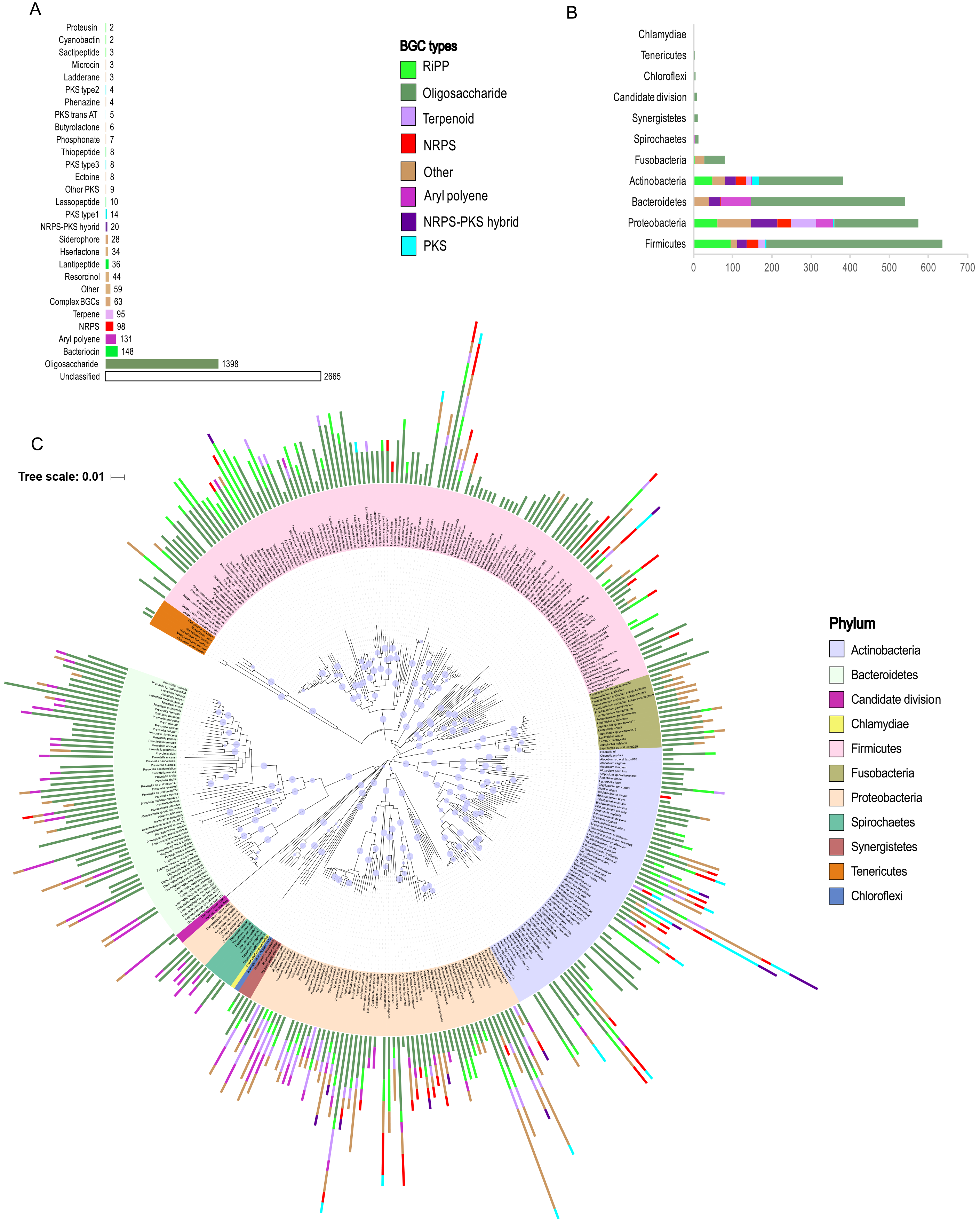
The oral microbiome contains a massive diversity of BGCs encoded by a multitude of taxa. **(A)** Bar graph illustrating the most common BGC subtypes identified in this study. Bars are colored according to higher level BGC class. **(B)** Bar graph illustrating the distribution of eight major classes of BGCs by phyla. **(C)** Phylogenetic tree based on 16S rRNA gene sequences showing the distribution of BGCs encoded by oral bacteria. Nodes with bootstrap values higher than 80% are displayed in the tree. Numbers of BGC types identified within each genome are shown in the bar graph and colored by BGC type. Leaf labels are colored by phyla. antiSMASH often identifies BGCs that encompass multiple gene clusters of different types fused into a single large gene cluster. 63 (~3%) of such unresolved BGCs and were encountered, and were categorized as the ‘complex’ BGC type (For convenience, we combined these BGCs with BGC types ‘Other’ for subsequent analysis). Distribution of BGCs is presented in more detail in Fig. S1.

Of the BGCs of a known class, a substantial fraction (1,398 BGCs, 62%) were annotated as oligosaccharides, making it the most abundant class of BGCs in the oral cavity. Oligosaccharide pathways are widely distributed across bacterial phyla and are predominant in Firmicutes, Proteobacteria, Bacteroidetes, Actinobacteria and Fusobacteria, with the highest number being identified in Firmicutes (Fig. 1B and C). Their ecological roles are largely underexplored, but studies show important functions such as capsule formation in virulence development (32) and attachment to surfaces, including neighboring bacterial species and host cells (33). Furthermore, diffusible oligosaccharides are known to display antibacterial activities (34), for example a previous study showed that polysaccharide A from the human gut bacterium *Bacteroides fragilis* can modulate the gut mucosal immune response (35, 36).

Another highly represented BGC class was ribosomally synthesized and post-translationally modified peptides (RiPPs), for which 209 BGCs (9.3% of BGCs of a known class) were identified. RiPPs include molecules such as bacteriocins, lantipeptides, sactipeptides, cyanobactins, and proteusins (denoted as fluorescent green in Fig 1). Of these RiPP types, bacteriocin-encoding BGCs were the most abundant as they contributed ~75% of the total RiPP diversity. Interestingly, although bacteriocin producing-BGCs were abundant in the oral microbiome overall, they were depleted in all Bacteroidetes genomes (Fig. 1C). The role of RiPPs, such as the bacteriocins, demands further exploration, as they exhibit antagonistic activities against other microbes sharing the same ecological niche, and influence competition for persistence between commensals and pathogens (37, 38). Furthermore, multiple studies genetic transformation in *Streptococcus* show that competence is tightly linked to bacteriocin production (39), which suggests that these molecules also play important roles in the horizontal transfer of genes and ultimately in niche differentiation and population structure changes.

BGCs encoding aryl polyene-like molecules in several Bacteroidetes and Proteobacteria genomes were identified (131 BGCs or 5.8% of BGCs of a known class). Aryl polyenes are predicted to function as protective agents against oxidative stress (40). However, only a few candidates have been experimentally characterized, leaving this group of small molecules highly underexplored. A diversity of non-ribosomal peptide synthetases (NRPSs), polyketide synthase (PKS), and NRPS-PKS hybrid BGCs (ranging between 0.9% and 4.4% of BGCs of a known class) were identified, in line with a prior study, which classified BGCs in the human microbiome in multiple body habitats (2). These compound classes are known for their antimicrobial activities and were previously characterized as possessing various nutrient-scavenging, immunosuppressant, surfactant, and cytotoxic properties (41). BGCs of the terpene class were also identified (95 BGCs, 4.2% of BGCs of a known class). This diverse group of small molecules may also be of ecological and medicinal interest since their activities have been reported as both anti-inflammatory and antimicrobial (42). The class ‘other’ encompasses BGCs that fall outside the known categories of antiSMASH-annotation, includes rare classes found in only few species, and constituted 9.4% of the total BGCs identified (Fig. 1A). Taken together, these results show that the oral microbiome encodes a vast and highly diverse array of small molecules that have largely unexplored, yet likely pivotal, roles in ecology and health.

### Sequence similarity networking reveals unexplored BGC diversity, even in well-studied classes of BGCs

In order to assess the evolutionary relationships between conserved domains in the proteins encoded by BGCs, as well as to group BGCs of similar putative function to evaluate novelty, a sequence similarity network approach was applied (see File S1). Briefly, the BGCs that were identified from the bacterial genomes using antiSMASH were aligned to the MIBiG repository (43) of 1,409 experimentally validated reference BGCs using the BiG-SCAPE algorithm (https://git.wageningenur.nl/medema-group/BiG-SCAPE). The resulting network comprised 4,242 nodes and 19,847 connecting edges revealing both close and distant homology to characterized biosynthetic pathways (Fig. 2). Notably, a significant fraction of the previously unclassified BGCs did sub-network with BGCs predicted to be of a known class, particularly the oligosaccharide, RiPP and aryl polyene classes (Fig. 2). This data provides inferences as to the function of over 100 previously unclassified novel BGCs.

**FIG 2:**
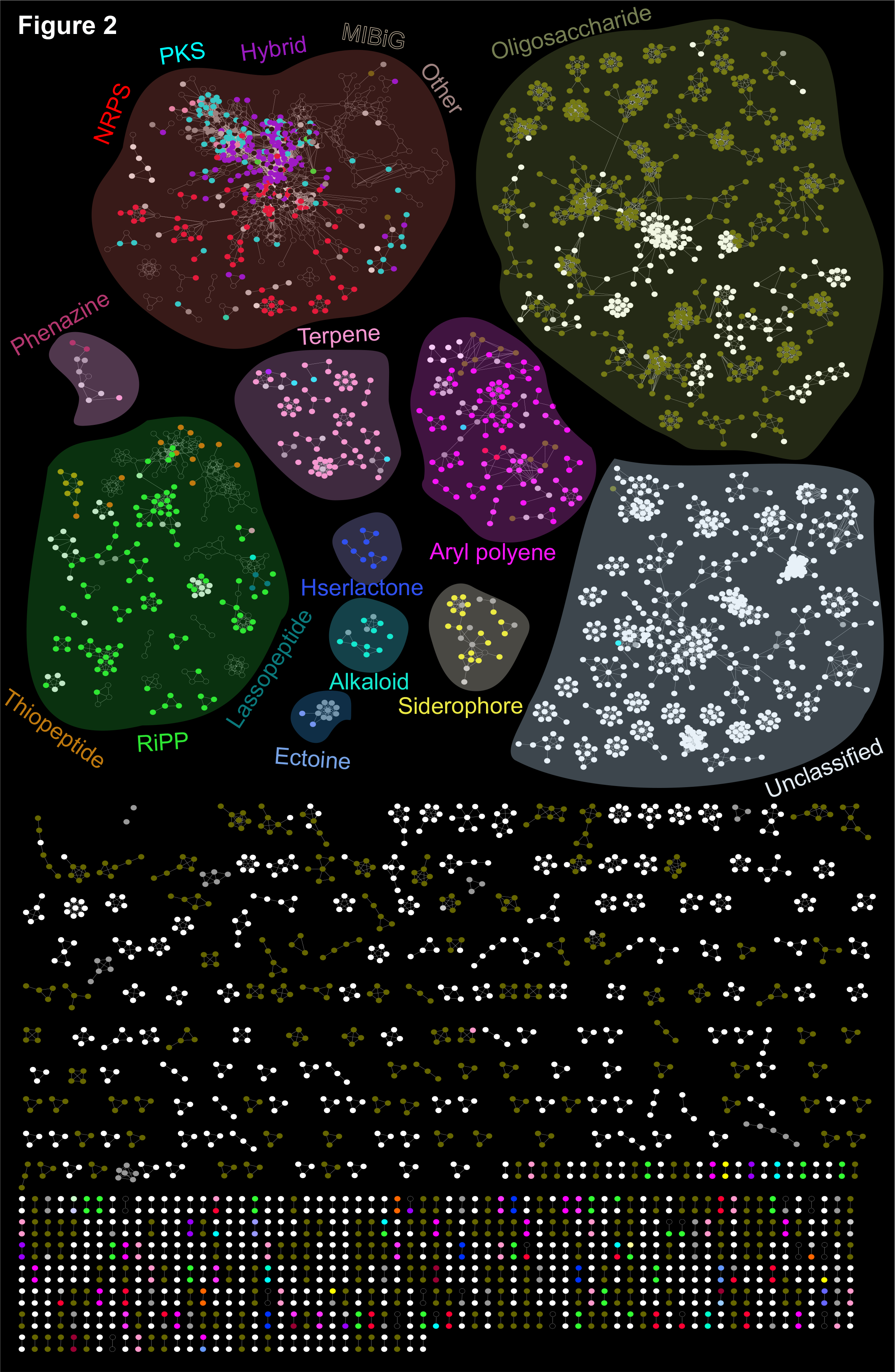
Similarity networking identified putative product classes for novel BGCs. Similarity network between the BGCs identified in the oral cavity and the experimentally characterized reference BGCs obtained from the MIBiG repository. Sub-networks representing major BGC classes, as determined by antiSMASH and BiG-SCAPE, are highlighted with different background colors to visualize BGCs as constellations within the biosynthetic landscape. Nodes (small circles) represent amino acid sequences of BGC domains and are colored by BGC class. Unfilled nodes represent reference BGCs from the MIBiG repository. Edges drawn between the nodes correspond to pairwise distances, computed by BiG-SCAPE as the weighted combination of the Jaccard, adjacency anddomain sequence similarity indices. For increased simplicity, only sub-clusters of unclassified and oligosaccharide BGCs with a minimum number of eight nodes are organized into given highlighted constellation.

The largest sub-network, comprised of mainly oligosaccharide-encoding BGCs, showed no significant homology with any experimentally validated BGCs in the MIBiG repository (Fig. 2). This may be due to the fact that oligosaccharide-producing BGCs are in at times categorized with primary metabolism, and not natural product-producing BGCs, as is the case in this study. The second-largest major sub-network was comprised of primarily unclassified BGCs. These may encompass distinct chemical scaffolds, and may represent a rich source of novel BGC types. The NRPS, PKS, NRPS-PKS hybrids, and a few terpene BGCs, grouped together forming a subnetwork implying a set of common core domains involved in these biosynthetic assembly lines, as described previously (4, 41). The majority of NRPS, PKS, NRPS-PKS hybrids, and RiPPs (in particular thiopeptides and lantipeptides) showed strong associations with MIBiG reference BGC sequences. It should be noted that these are the most prevalent classes in the MIBiG repository (Table S2). Currently, only four experimentally characterized aryl polyene BGCs exist in the MIBiG database, therefore it was not surprising that none of the nodes in the aryl polyene cluster sub-networked with MIBiG reference BGCs. Given that aryl polyenes are thought to be the most abundant BGC class in the human microbiome (4), this indicates that this class of molecules is severely understudied (Fig. 1 and Table S2). Several BGCs annotated as saccharides, other, unclassified, PKS and NRPS BGC types grouped with aryl polyene BGCs, which may represent novel hybrid classes of BGC. Other small sub-networks include biosynthesis of terpene phenazine, homoserine lactone, alkaloid, siderophore, and ectoine. These sub-networks did not associate with MIBiG reference BGCs, indicating that they also await experimental validation. Our implemented analysis approach, using the MIBiG/BiG-SCAPE pipeline, is powerful with regards to predicting the functions of novel BGCs. The annotations we generated here provide deeper insights of which BGCs and compound classes are most likely to be identified in futures studies, due to knowledge of their closest neighbor’s biochemical properties. The BGCs remaining with completely unknown functions represent exciting future challenges, which could be addressed by generating large-insert BGC expression libraries.

While antiSMASH and network analysis were employed for broad classification of BGCs into known classes, MultiGeneBlast was also utilized at the level of the entire gene cluster to further annotate BGCs in-depth and identify homologs against the MIBiG repository (30). Using this approach, the 4,915 BGCs were classified into four major categories based upon the level of homology to known experimentally validated BGCs in the MIBiG repository. This categorization resulted in 1,146 (20%) BGCs closely homologous, 848 (15%) BGCs moderately homologous and 2,221 (40%) BGCs distantly homologous to well-characterized BGCs (Fig. S2). 1,393 (~25%) BGCs did not appear to have significant homology to BGCs in MIBiG, based upon the E-value (see Methods section for details). Such a detailed annotation of BGCs harbored by the human oral microbiome has not been accomplished previously.

### Specific BGCs are associated with periodontitis and dental caries

We next systematically examined the differential representation of bacterial BGCs in saliva and dental plaque across 294 human subjects with good oral health, dental caries, or periodontitis. The data from 247 subjects was obtained from eight previous studies, which represented all publicly available metagenomes and metatranscriptomes associated with caries or periodontal disease, compared to health, at the time of this study (Table S3). In addition, DNA from 47 saliva samples representing 23 children with caries and 24 healthy children was sequenced and putative BGCs were identified (see Fig S3 for workflow). Non-supervised exploratory ordination through PCoA revealed significant differences in the representation of BGCs between healthy and diseased subjects in five of the six metatranscriptome studies and six of the seven metagenome studies investigated (Fig. S4). The 1,804 BGCs which were differentially represented in health versus disease in the metagenomes and metatranscriptomes are summarized in Table 1.

**Table 1.** Number of differentially represented or expressed biosynthetic pathways in saliva, supra- and sub-gingival plaque samples, from shotgun metatranscriptomics and metagenomics libraries representing 231 subjects (oral health: n = 110, dental caries: n = 77, periodontitis: n = 44). 741 BGCs were differentially abundant in caries (515 enriched, and 226 less abundant) and 1,063 BGCs were periodontitis associated (670 enriched, and 393 less abundant). 355 BGCs were differentially expressed in caries (208 up- regulated, and 147 down-regulated), while 421 BGCs were either up- or down-regulated in subjects with periodontitis (218 up-regulated, and 203 down-regulated).

| BGC type | Metatranscriptome (up/down) |  | Metagenome (up/down) |  |
| --- | --- | --- | --- | --- |
|  | Caries | Periodontitis | Caries | Periodontitis |
| Aryl polyene | 4/11 | 29/4 | 26/5 | 25/3 |
| Bacteriocin | 14/10 | 1/1 | 31/1 | 5/24 |
| Butyrolactone | 2/0 | 1/1 | 3/0 | 1/1 |
| Homoserine lactone | 0/4 | 3/4 | 10/0 | 1/1 |
| Lantipeptide | 7/0 | 4/0 | 11/2 | 1/6 |
| Lasso peptide | 0 | 0 | 0 | 0/3 |
| NRPS | 7/0 | 2/8 | 11/7 | 10/14 |
| NRPS-PKS hybrid | 1/0 | 0/1 | 5/0 | 0/1 |
| Oligosaccharide | 66/68 | 100/40 | 116/118 | 287/132 |
| Other | 7/0 | 2/11 | 36/3 | 16/15 |
| Phenazine | 0 | 0/1 | 2/0 | 0/1 |
| PKS | 3/0 | 1/0 | 8/0 | 1/4 |
| Proteusin | 0 | 0 | 1/0 | 1/0 |
| Resorcinol | 1/4 | 3/8 | 4/4 | 19/0 |
| Sactipeptide | 0 | 0 | 0 | 1/0 |
| Siderophore | 1/0 | 1/2 | 0 | 1/0 |
| Terpene | 1/10 | 9/1 | 42/0 | 1/4 |
| Thiopeptide | 1/0 | 0 | 0 | 0/7 |
| Unclassified | 93/50 | 62/121 | 209/86 | 300/177 |

The BGCs associated with disease in the metatranscriptome studies were related to the synthesis of a broad range of small molecule types. These particularly included BGCs of the oligosaccharide, aryl polyene, terpene, bacteriocin and NRPS classes (Fig. 3). BGCs encoding PKS, NRPS, and bacteriocins from *Actinomyces, Rothia* and *Corynebacterium* had increased expression in subjects with caries, while BGCs encoding terpenes and aryl polyenes from *Neisseria spp.* and Proteobacteria had increased expression in healthy subjects (Fig. 3). Previous studies illustrated that aryl polyenes act as protective agents against oxidative stress, and that terpenes function as antiinflammatory agents (40). Interestingly, high levels of *Actinomyces* were previously associated with severe early childhood caries (44). In the caries associated samples, known caries-associated species belonging to the *Streptococcus, Veillonella*, and *Lactobacillus* genera (45) showed notable changes in bacteriocins and oligosaccharides BGC expression profiles (Fig. 3).

**FIG 3:**
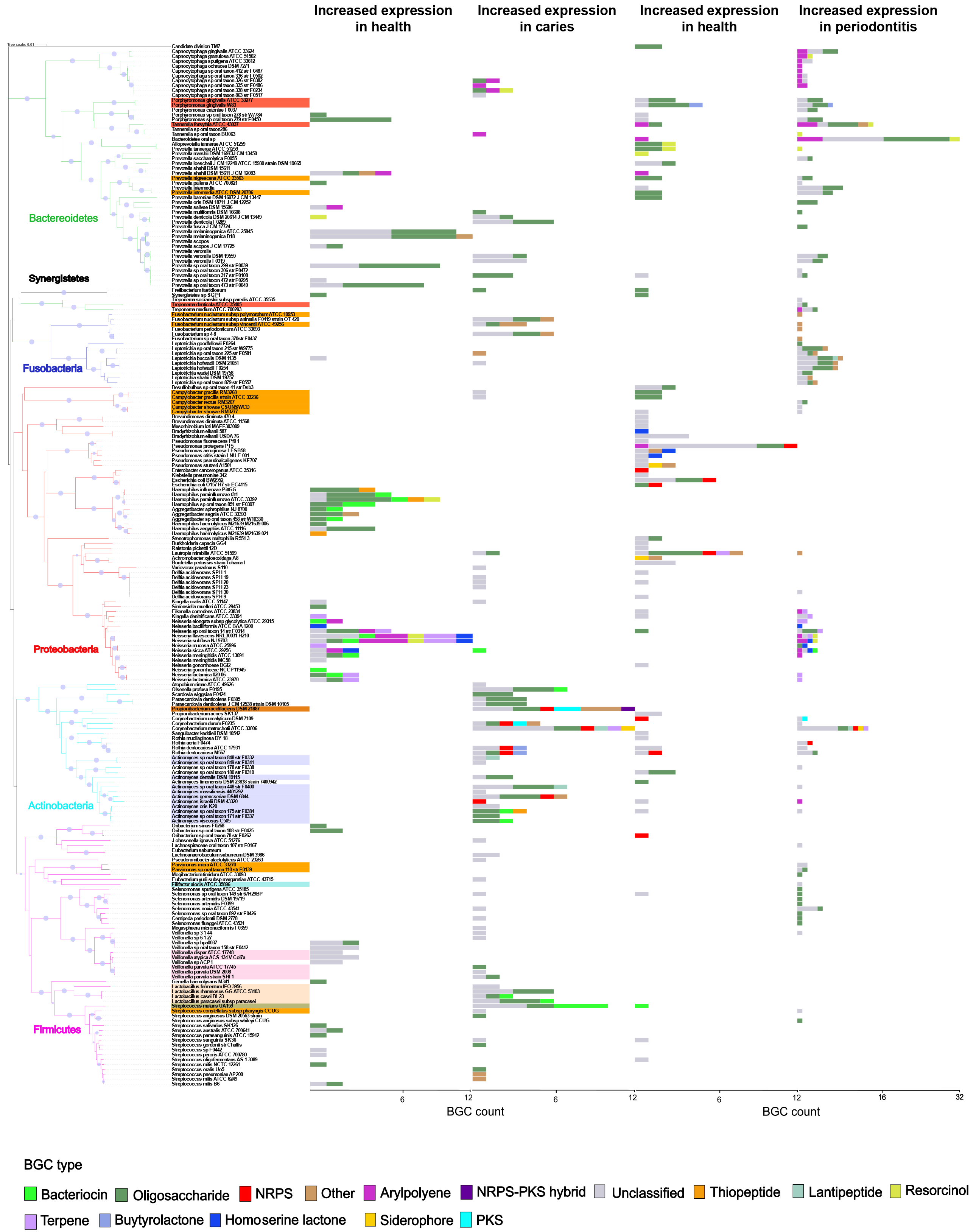
BGCs are differentially expressed in health and disease. Bar graphs illustrating phylogenetic distribution of biosynthetic pathways in health- and disease-associated oral microorganisms. Species with significant changes in BGC expression based on the analyzed metatranscriptomic data sets are shown in the phylogenetic tree on the left. Bar graphs at the leaf tips display number of BGCs either over or under expressed and colored according to the BGC type. It should be noted that the x-axis scales are different in left and right panels. Significant differences in the expression of BGCs were determined based on negative binomial distribution model using DESeq2 with FDR correction (p-value < 0.05).

In periodontitis, a high number of differentially expressed BGCs (170 BGCs) were identified in community members belonging to the Bacteroidetes phylum. Interestingly, several BGCs encoded by periodontal pathogens of the red and orange complexes (e.g. *Porphyromonas gingivalis*) were differentially expressed in health compared to periodontal disease. Known red complex species had increased expression of BGCs belonging to the aryl polyene, oligosaccharide, homoserine lactone and resorcinol classes in diseased states. *Neisseria spp.* also showed interesting signatures, such as increased expression of BGCs belonging to the terpene, resorcinol, bacteriocin, and homoserine lactone classes (Fig. 3). Homologs to specific BGC products in the MIBiG database which displayed differential expression in health and disease are detailed in Figure S5. Analysis of the metagenomic studies yielded similar trends to those detailed above (Figs. S6 and Fig. S7).

Next, a subset of differentially represented BGCs, which showed high expression in either healthy or diseased states, was examined to determine if they commonly occur across studies. The results were visualized as a binary occurrence matrix (Fig. S8). In all studies analyzed, only a minor fraction of the differential features (< 10 BGCs) were shared between any two studies. Besides high inter- and intra-individual variations in the microbial composition, the significant study-to-study variation can likely be attributed to differences in sequencing platforms (Table S3). This factor may have influenced the sequence composition and sequencing depth, particularly considering the metagenome and metatranscriptome complexity (Fig. S9 and Table S4). Based on the above comparisons, the authors suggest that differences between sequencing and computational platforms (e.g. alignment parameters and sequence read filtering) must be considered, and that future efforts to obtain high-quality, deep-coverage sequencing data will help alleviate the study-to-study noise observed here.

### Correlations between BGCs and oral taxa are depleted in dental caries as compared to health

To examine the relationship between BGCs and bacterial taxa during health and disease, a focused comparative analysis of the shotgun metagenomics data obtained in this study from healthy children and children with caries was performed. Interactions between BGCs and microbial taxa were examined by employing cooccurrence network analysis using the SparCC algorithm, which has the benefit of limiting the number of spurious correlations identified due to species data being compositional (46). While positive correlations were more evident among taxa-taxa relationships, (i.e. different taxa benefit from one another’s presence), almost all significant correlations that were identified between specific BGCs and taxa were negative (Table S5 and Figs. S10 and S11). This suggests that antagonistic relationships, modulated through BGC-produced antimicrobial molecules, are highly significant to the ecology of the oral microbiome.

All BGCs which had significant correlations to oral taxa (a total number of 36) were annotated as close homologs to previously characterized BGCs belonging to the PKS, NRPS, NRPS-PKS hybrid, oligosaccharide and aryl polyene classes (Fig. S10). In the oral microbiomes derived from healthy children, the interaction network was dominated by negative correlations between oral taxa and BGCs producing glycopeptidolipids, capsular polylsaccharides, as well as a homolog of flexirubin (Fig. S11A). The glycopeptidolipids were encoded by the opportunistic pathogens *Kytococcus sedentarius* and *Mycobacterium neoaurum*, and were primarily shown to vary inversely with the oral taxa *Lactobacillus, Prevotella, Capnocytophaga* and *Enterococcus* (Fig. S11A and Table S5). The flexirubin homolog BGCs were encoded by *Actinomyces massiliensis* and *Prevotella oralis*, and displayed antagonistic activity against 122 taxa, including *Streptococcus mutans*, historically considered the primary etiologic species of dental caries (Fig. S11A and Table S5). Homologs of the antibiotics bacillaene and pristinamycin (47, 48), harbored by genomes of the health-associated species *Propionibacterium propionicum* F0230a and *Actinomyces timonensis* DSM 23838 (Fig. S11A, Table S5), displayed negative correlations with several pathogenic taxa: *Lactobacillus, Listeria, Lysinabacillus, Acinetobacter, Enterococcus, Neisseria, Staphylococcus, Kingella* and *S. mutans* (49) (Table S5). These associations are reminiscent of a previous study which observed similar macrolide-encoding BGCs widely distributed amongst oral bacterial genomes (2). These macrolide structures were also reported to inhibit the growth of cariogenic Streptococci (50). This collective evidence indicates that the isolation and characterization of bacillaene- and pristinamycin-like molecules in future studies may be key to understanding important health-protective mechanisms in the oral cavity. Finally, *P. propionicum* F0230a encoded a BGC with high sequence homology to a non-ribosomal peptide pathway encoding the genotoxin, colibactin (51). This BGC showed antagonistic associations with pathogenic genera: *Haemophilus, Aggregatibacter, Parascardovia, Capnocytophaga* and *Streptococcus*.

Most intriguingly, the number of significant correlations between BGCs and microbial taxa was dramatically reduced in the samples derived from children with caries (Fig. S10A to C). This may indicate that in the oral cavities exhibiting disease, the well-documented colonization resistance of the oral microbiome may be impaired. Of the few significant correlations between BGCs and taxa within the interaction network of the caries-associated microbiome, the vast majority involved BGCs encoding RiPPs with close homology to nosiheptide and hygromycin BGCs (Fig. S11B, Table S5). The nosiheptide-like BGC, encoded in the genome of *C. matruchotii*, was the most predominant, with antagonistic interactions against ~90 taxa. These included pathogens from the *Klebsiella*, *Helicobacter*, *Filifactor*, *Haemophilus*, *Enterococcus*, *Fusobacterium* genera (Fig. S11B, Table S5). The hygromycin-like BGC from *P. propionicum* negatively correlated with several pathogens belonging to the genera *Lactobacillus, Neisseria, Klebsiella, Anaerococcus* and *Pseudoramibacter*. Interestingly, there were no significant correlations between *S. mutans* and BGCs in the caries-associated oral microbiomes, which may indicate that during disease, the community lacks the ability to limit the abundance of this keystone pathogen. Taken together, these results suggest that in the oral microbiome, exclusion of particular taxa via antagonistic interactions, mediated by the products of BGCs, is widespread (Table S5). Although such interactions were still present in the caries-associated oral microbiomes, they were much fewer in number. This underscores the importance of ecology, and the role of BGC-produced small molecules, in the balance between health and disease.

### Homologs of BGC-produced small molecules are present in oral metabolomes associated with caries and health

To validate the production of small molecules by differentially abundant BGCs, untargeted liquid chromatography-tandem mass spectrometry (LC-MS/MS) analysis of saliva samples was performed. Utilizing the Global Natural Products Social Molecular Networking (GNPS) (52) analysis platform, a mass spectral molecular network consisting of 1,369 mass spectral features grouped into 69 molecular families (two or more connected components of a graph) was obtained. 50 matches were acquired between the query MS/MS spectra and characterized reference spectra from GNPS. To further enhance mass spectrometry annotations and to link annotations to known chemical structures encoded by BGCs, major chemical classes were putatively identified by integrating mass spectral molecular networking with *in silico* annotations and automated chemical classification approaches (53–55). This allowed identification of approximately 38% of the nodes in the mass spectral molecular network at the chemical class level. The most predominant chemical classes within the network were carboxylic acids and derivatives, prenol lipids, fatty acyls, and flavonoids (Fig. S12). Substructures associated with macrolides, terpenoids, and macrolactams were also identified. At the chemical class level, distinct relative abundance patterns between the health and disease-associated samples could be observed for carboxylic acids and derivatives. The PCoA analysis of the 1,369 unidentified MS features, showed clear separation of samples between healthy and diseased states (Fig. S13A), in agreement with the BGC abundance profiles (Fig. S4M). By employing a random forest importance model, 15 key metabolites, which were distinct between healthy and disease states (Fig. S13B), were identified. 12 of the 15 key metabolites were significantly more abundant in healthy subjects, while three were more abundant in the subjects with dental caries. Out of the three key metabolites that were significantly more abundant in the diseased subjects, two matches were obtained to lipid compounds from GNPS reference spectra resulting in a level-2 metabolite identification (56). These matches were N-Nervonoyl-D-erythro-sphingophosphorylcholine and 13-Docosenamide. These molecules are likely to originate from the human host and warrant further investigation.

Using the *in silico* Network Annotation Propagation tool (NAP) (57), putative structural matches were obtained for 6 out of the 12 key metabolites that were more abundant in the healthy subjects, including terpenoids, phenylpropanoids as well as fatty alcohols. It should be noted however, that one of the limitations of *in silico* annotation is the uncertainty around the correct structure among the predicted candidate structures. Results should therefore be interpreted with care, and an accurate prediction of the putative identity would require follow-up investigations, which is outside the scope of the present study. It should be also noted that both the genomics and metabolomics approaches employed identify putative homologs and not exact matches. Thus, using current techniques and databases, is it not possible to definitively determine if the small molecules identified by LC-MS/MS were in-fact produced by the specific BGCs predicted by antiSMASH. However, the LC-MS/MS analyses largely support the results of the genomic analyses by detecting classes of small molecules and homologs which were similar to those discovered by the complementary BGC genomics analyses.

### Concluding remarks

This study significantly expands the number of identified BGCs encoded by bacteria of the human oral microbiome and designates putative products to many novel clusters. Representation and expression of the newly identified BGCs, as well as their relationship to the abundance of oral bacterial taxa was examined during health, dental caries, and periodontitis, revealing significant differences in microbial social ecology and communication among the three host outcomes. This work provides an atlas for further examination and experimental validation of the identified socio-chemical relationships and their role in the pathogenesis of dental caries and periodontal disease. A deeper elucidation of the social activities of the microbes residing in the oral cavity will significantly improve our understanding of the pathogenesis of oral (and extra-oral) diseases and will guide development of improved therapeutic strategies to maintain health.

## MATERIALS AND METHODS

The ethics statement is provided in the Supplementary Materials and Methods section of Supplementary Material File S1

### Study inclusion/exclusion criteria and collection of saliva

Approximately 2 ml saliva was collected by spitting method in a 15 ml Falcon tube over a 20 min period. Whole saliva was immediately transferred to sterile 2 ml cryovial tubes and centrifuged at 6000 × g for 5 minutes to remove eukaryotic cells and solid debris. Supernatants were collected, mixed with glycerol (20%), and snap-frozen for long term storage at −80°C. For detailed protocol, see Supplementary Materials and Methods.

### DNA extraction and metagenomics sequencing

For a detailed protocol, see Supplementary Materials and Methods.

### BGC identification and network analysis of known and putative oral BGCs

A list of 1,362 described and curated human oral taxa (18^th^ September 2017) was obtained from HOMD, Human Oral Microbiome Database (55). In order to identify small molecule and secondary metabolite-encoding BGCs in genomes of bacterial taxa representative of a broad oral bacterial diversity, 461 complete and high-quality draft genomic sequences, annotated as dynamic and static, were obtained from the National Center of Biotechnology Information genome database (http://www.ncbi.nlm.nih.gov/genome), as well as from an in-house database (Table S1). These were concatenated into a major query-database and fed to antiSMASH, (Antibiotics & Secondary Metabolite Analysis Shell, version 4.0) (29). Multiple nucleotide FASTA sequences from BGCs were constructed. We excluded a list of 320 previously described non-biosynthetic genes commonly found in BGCs (2) (Table S6) based on text within an attribute using advanced filter settings in CLC Workbench software v. 9. (CLCbio, Aahus, Denmark). The resulting dataset contained a total of 192,283 gene sequences from 4,915 BGCs and can be downloaded from the MassIVE repository (https://massive.ucsd.edu/) with the accession ID MSV000081832. For more information, see Supplementary Materials and Methods.

### Comparison of BGCs with known biosynthetic pathways

A reference MIBiG database comprising multiple amino acid sequences for each BGC was constructed using MultiGeneBlast (30). To further compare BGCs derived (excluding the fatty acid synthase encoding BGCs) from oral bacterial genomes with those encoding the biosynthetic pathways for known compounds, we performed multi-gene homology searches using complete gene cluster sequences against the MIBiG database by using the stand-alone version of MultiGeneBlast (http://multigeneblast.sourceforge.net/) algorithm with default settings. Subsequently, for each queried BGC, we extracted information from the top hit (with the highest cumulative BLAST bit score) from an output of multiple BLAST hits using an in-house python script. For additional information, see Supplementary Materials and Methods.

### 16S rRNA gene (16S) phylogenetic analysis

For a detailed protocol, see Supplementary Materials and Methods.

### Metagenomic and metatranscriptomic data collection

Shotgun metatranscriptomic and metagenomic sequencing data published previously by Duran-Pinedo et al. (22), Belda-Ferre et al. (23), Belstrøm et al. (24), Jorth et al. (58), Do et al. (25), Peterson et al. (26), Yost et al. (27), Wang et al. (28), and Shi et al. (59), as well as our own study of metagenomes from saliva obtained from children with good dental health, or children with dental caries was analyzed (sequence reads are accessible under BioProject PRJNA1234. Table S3). For detailed protocol, see Supplementary Materials and Methods.

### Differential abundance and expression analyses of BGCs

We employed a systematic workflow for analyzing abundance and expression profiles of the BGCs (see Fig. S3). Using SRA toolkit utilities, reads were extracted from metatranscriptome and metagenome shotgun sequenced libraries available via NCBI. For a detailed protocol, see Supplementary Materials and Methods.

### Principal Coordinate analysis

The differences between samples from healthy versus diseased individuals was investigated by applying Principal Coordinates Analysis (PCoA) on Manhattan distances generated on the DESeq2 normalized count file using the EMPeror (60) tool. For a detailed protocol, see Supplementary Materials and Methods.

### Correlation network analysis

The correlation network was constructed using the SparCC algorithm (46) python package (available at https://bitbucket.org/yonatanf/sparcc) to represent both co-abundance and co-exclusion networks between species and corresponding BGCs. For a detailed protocol, see Supplementary Materials and Methods.

### Experimental small molecule metabolites detection

Approximately 150μl of saliva was lyophilized and ethyl acetate was added to extract non-polar molecules. Samples were then vortexed, centrifuged to remove the cell debris and submitted to untargetd LC-MS/MS analysis. For a detailed protocol, see Supplementary Materials and Methods.

### Mass spectral molecular networking

LC-MS/MS spectra were preprocessed for feature extraction using MZmine2 (61) and submitted to mass spectral molecular networking through GNPS (43). For a detailed protocol, see Supporting Information.

### Putative chemical structure annotation

To putatively annotate chemical structures in our mass spectral molecular networks, we performed *in silico* structure annotation through Network Annotation Propagation (NAP) (57) both for [M+H]+ and [M+Na]+ adducts. For a detailed protocol, see Supporting Information.

## Acknowledgements

The authors thank Alexander Richter for providing custom python scripts to support the data analysis and thank Ricardo R. da Silva for supporting mass spectral analysis. This research was funded by NIH/NIDCR grants R00-DE024543-3 (A.E.), K99-DE024543 (A.E.), and F32-DE026947 (J.L.B.).

## Author contributions

G.A., X.T., C.Z. and A.E. designed the experiments, G.A., A.V.M., R.A., M.B.D., N.C.T. collected and processed the samples, G.A., X.T., C.Z. A.V.M. M.E. and A.E. analyzed the data, and G.A., J.L.B and A.E. wrote the manuscript. All authors helped edit the manuscript. All authors read and approved the final manuscript.

## Additional information

Correspondence and requests for materials should be addressed to A.E.

## Availability of data and material

Sequence data has been submitted to NCBI under BioProject ID PRJNA478018 with SRA accession SRP151559. Mass spectral files, LCMS/MS metadata file, Nucleotide FASTA sequences of the oral biosynthetic gene cluster collection are accessible from the MassIVE repository (https://massive.ucsd.edu/) with the accession ID MSV000081832.

---

**Figure S1.**
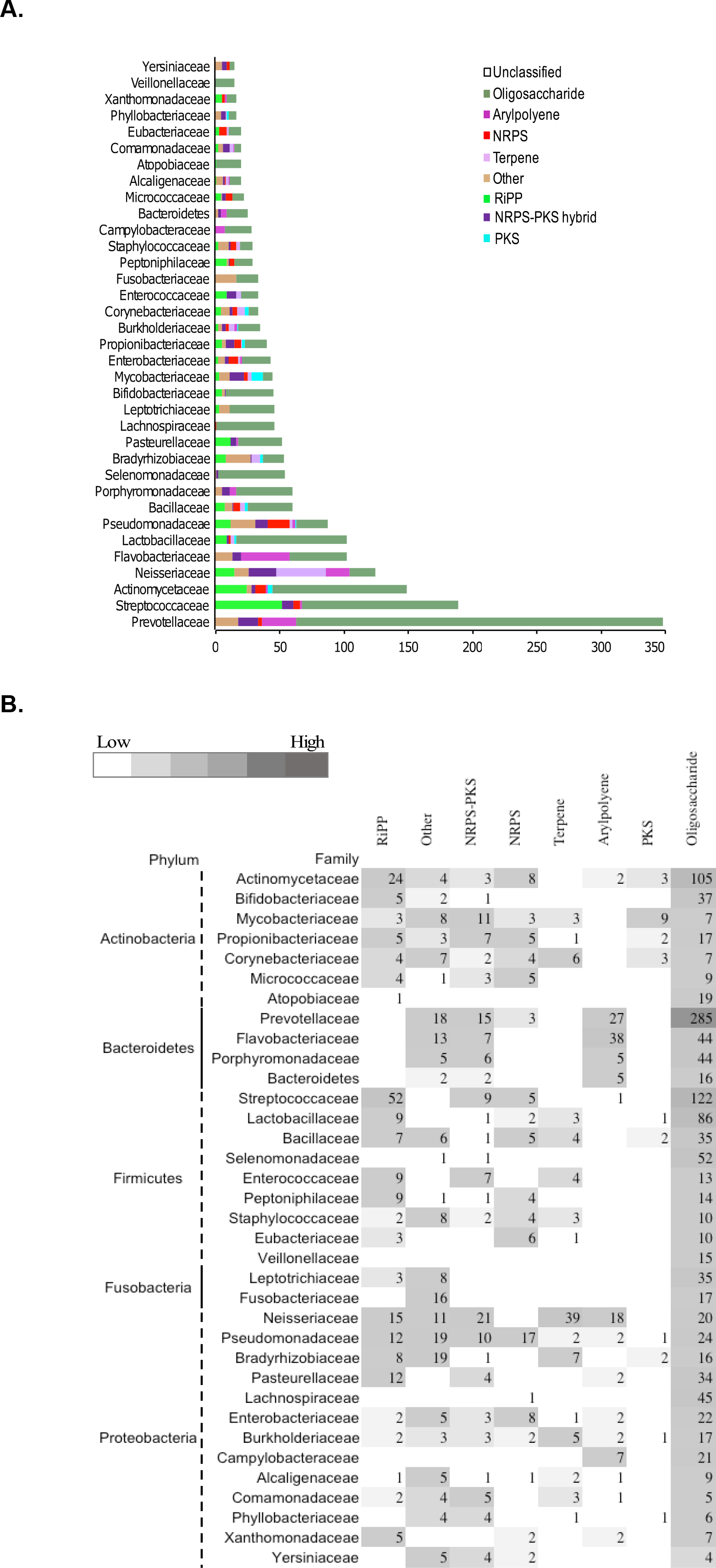

**Figure S2.**
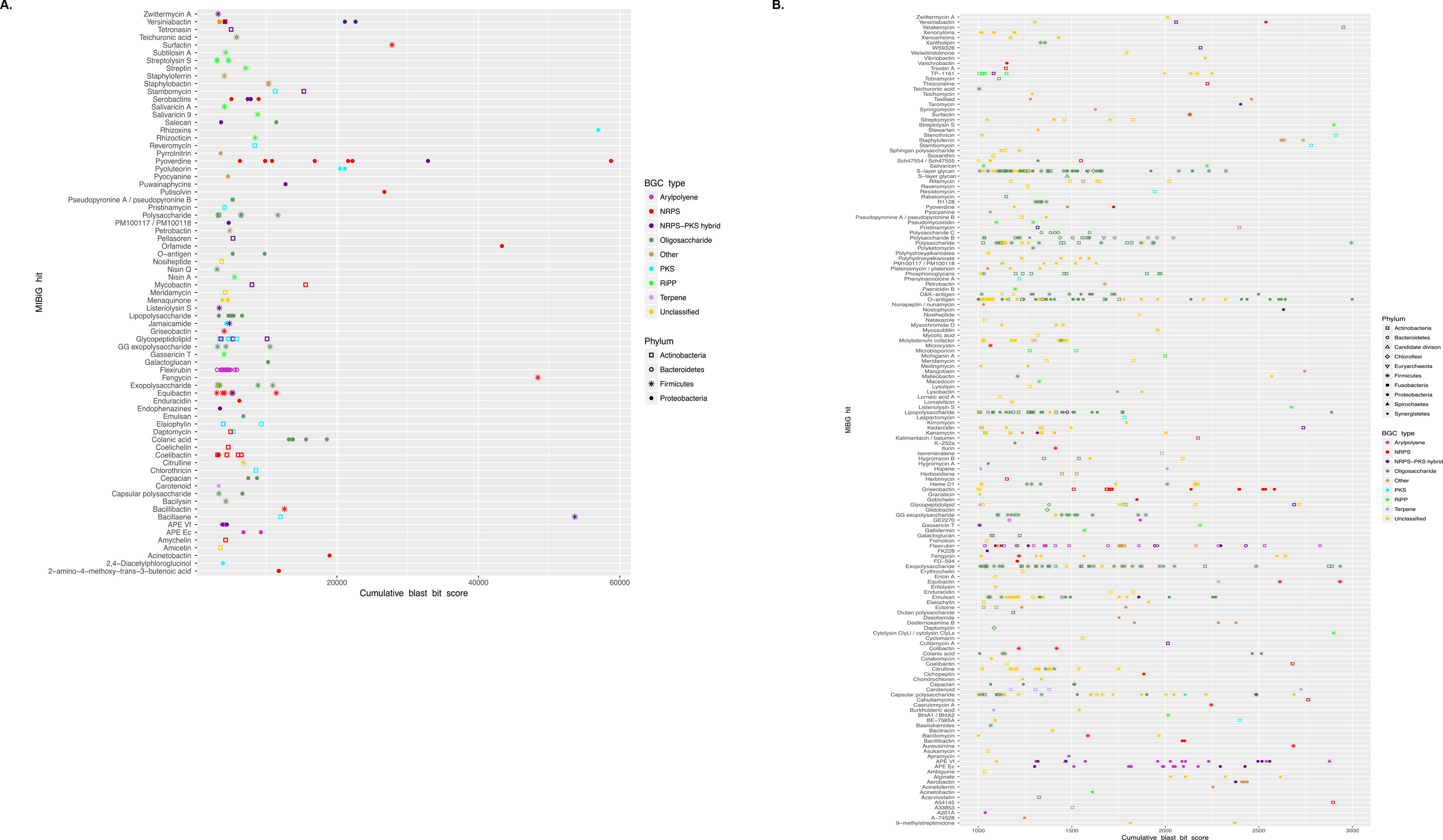

**Figure S3.**
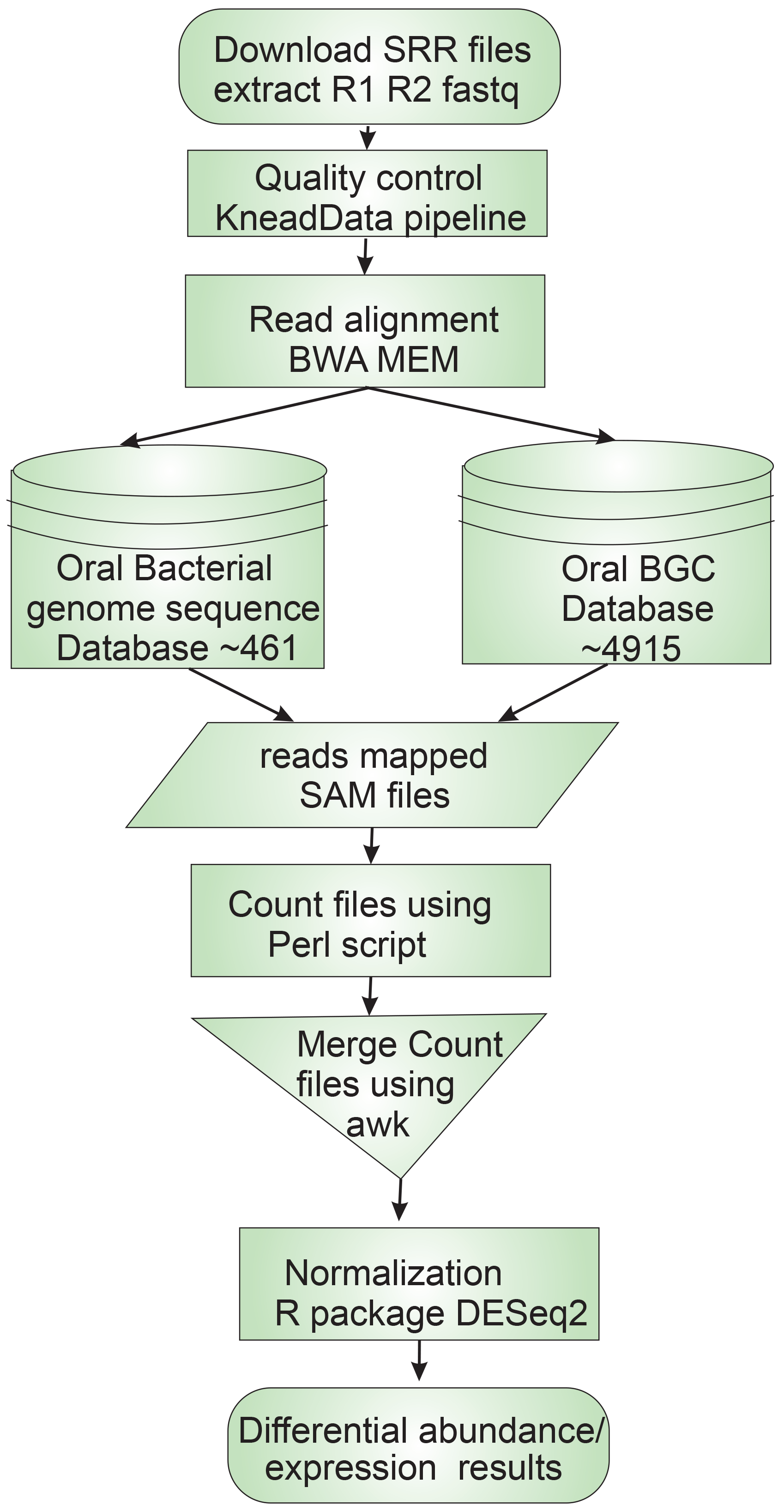

**Figure S4.**
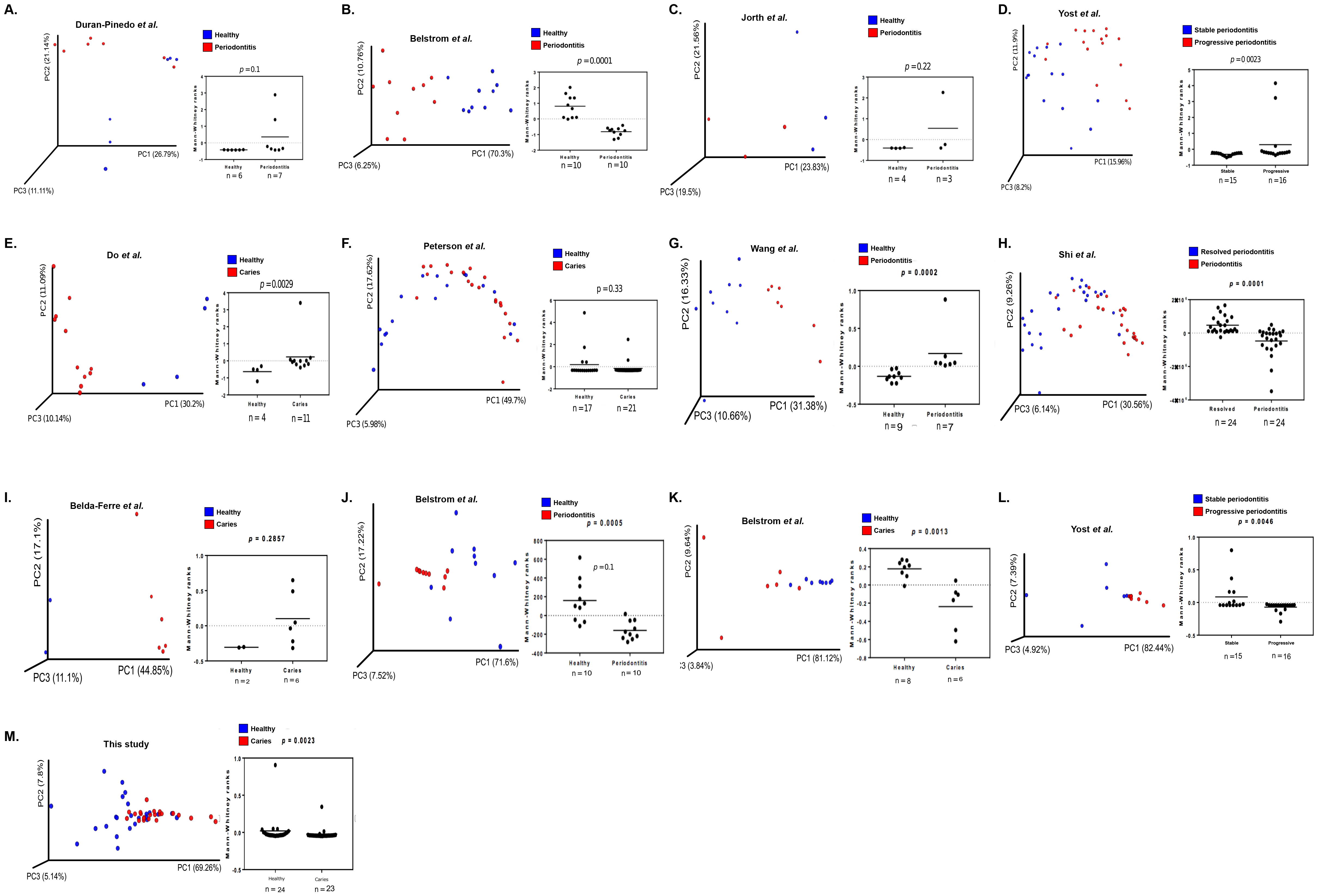

**Figure S5.**
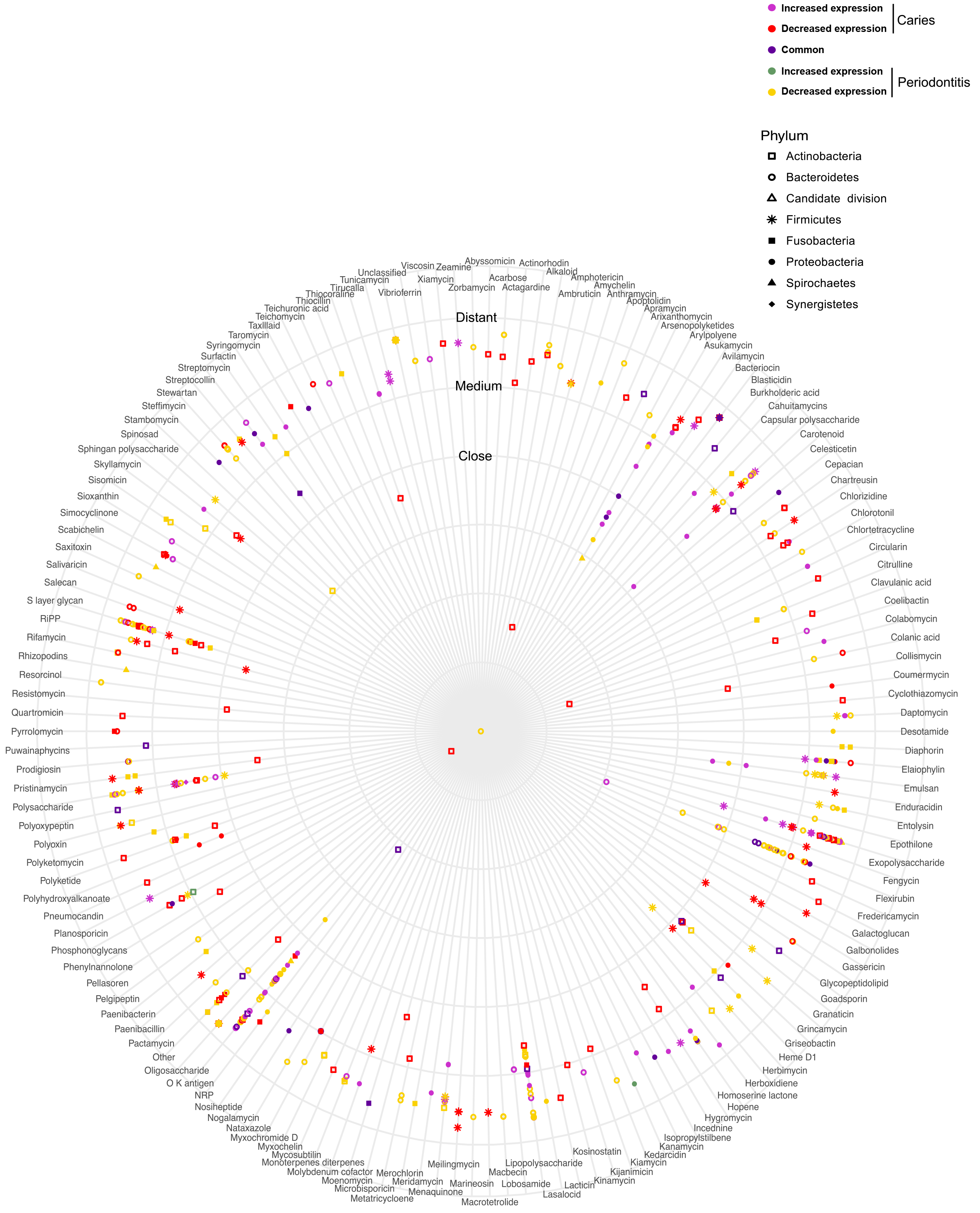

**Figure S6.**
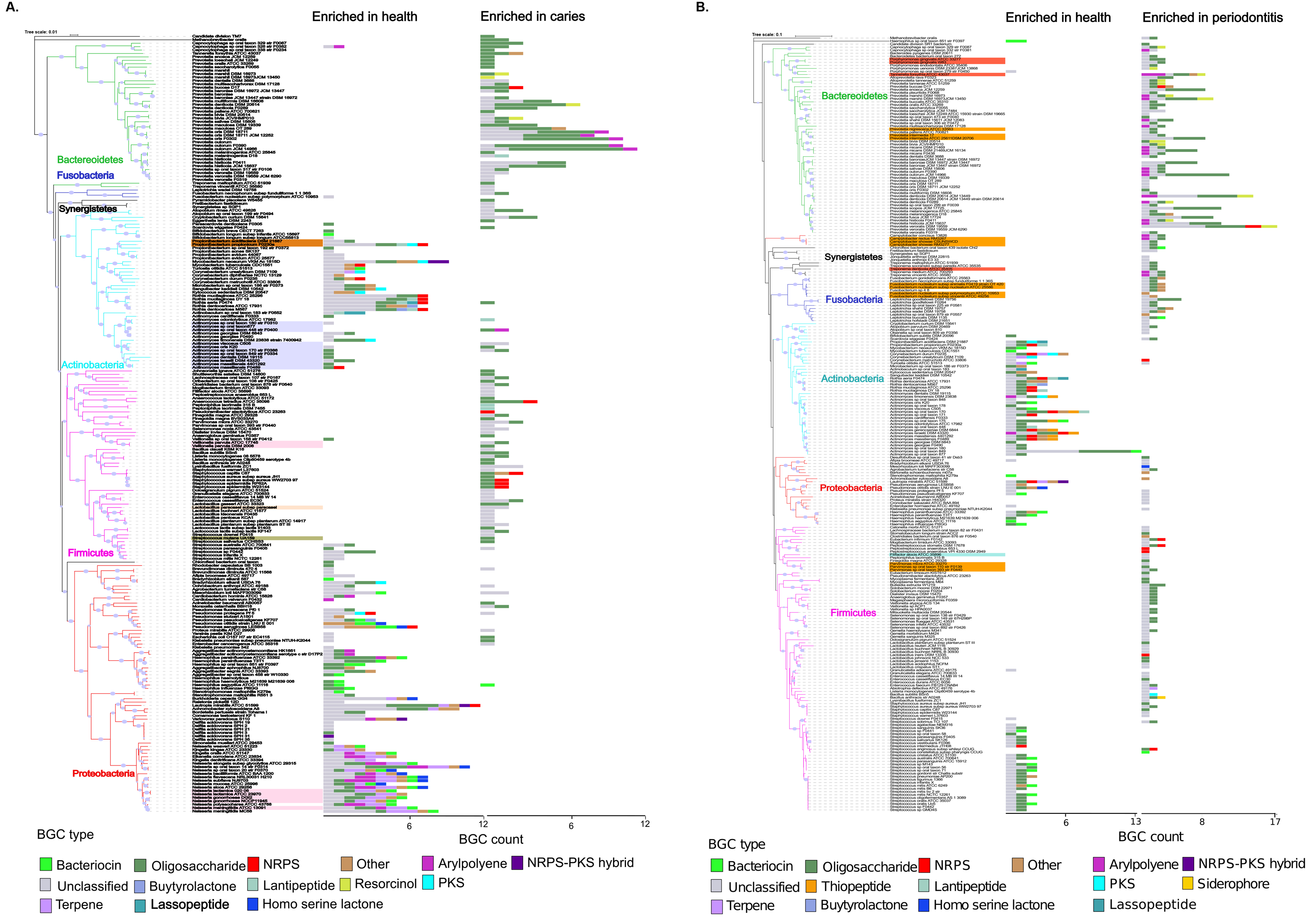

**Figure S7.**
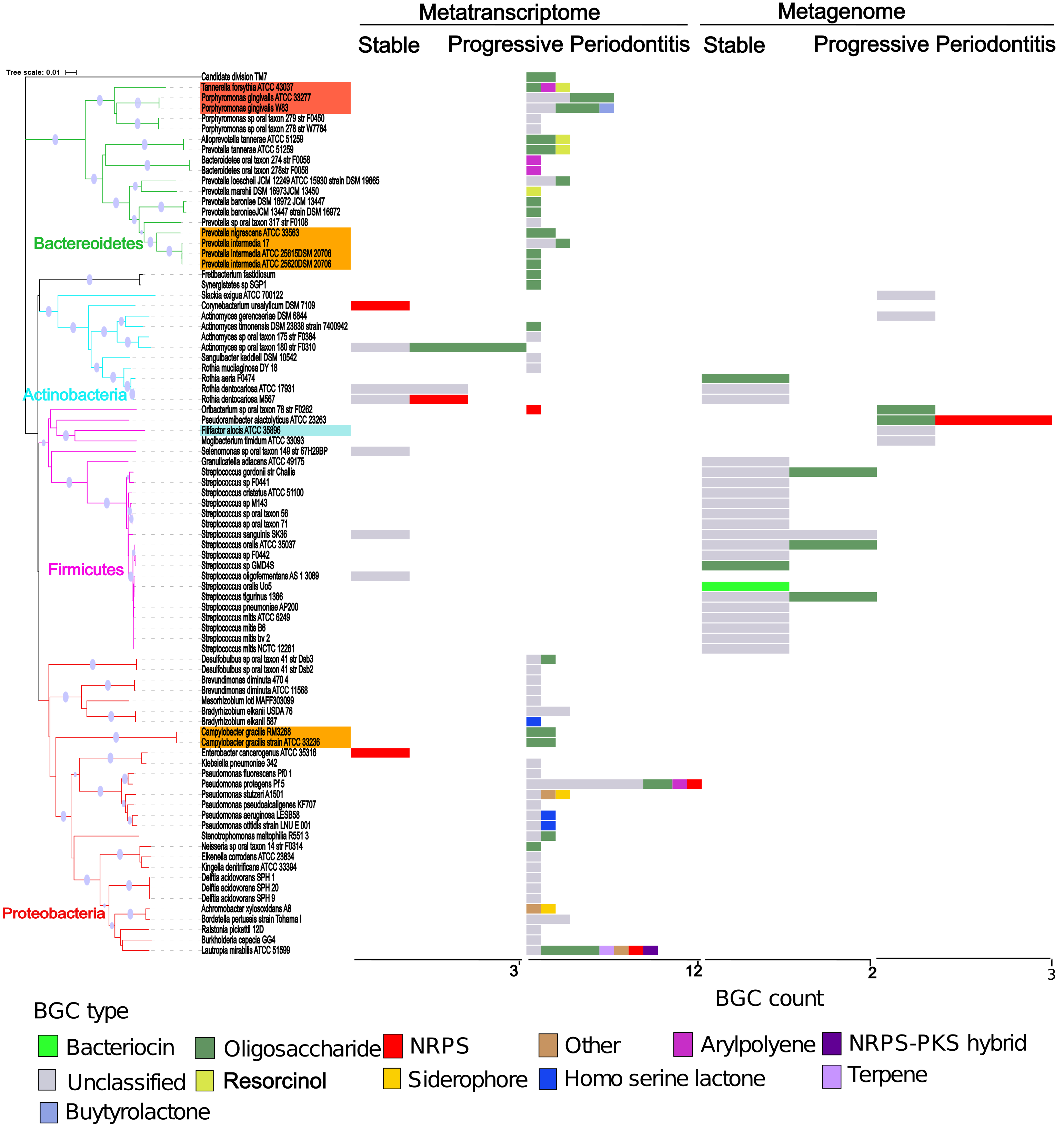

**Figure S8.**
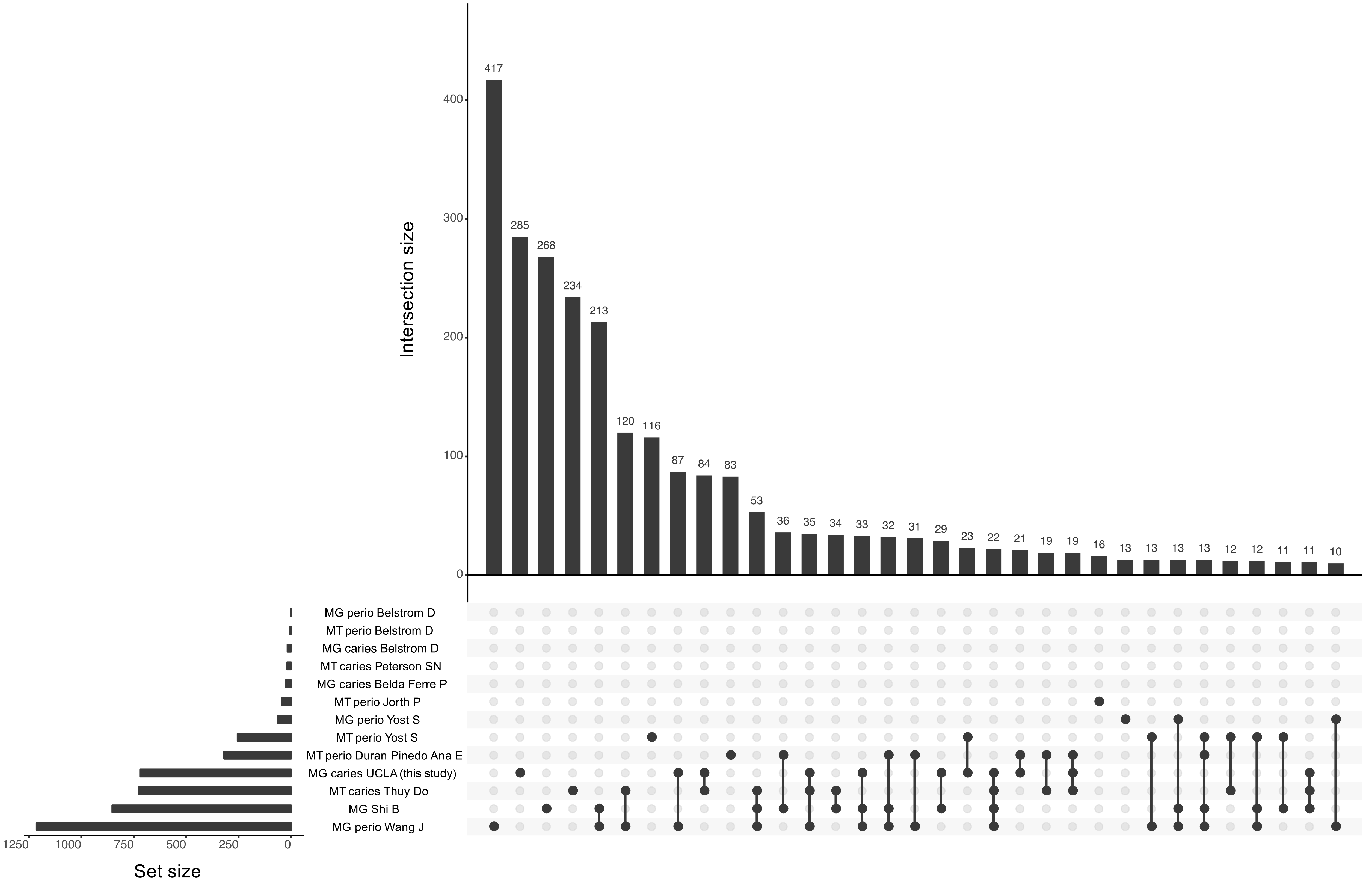

**Figure S9.**
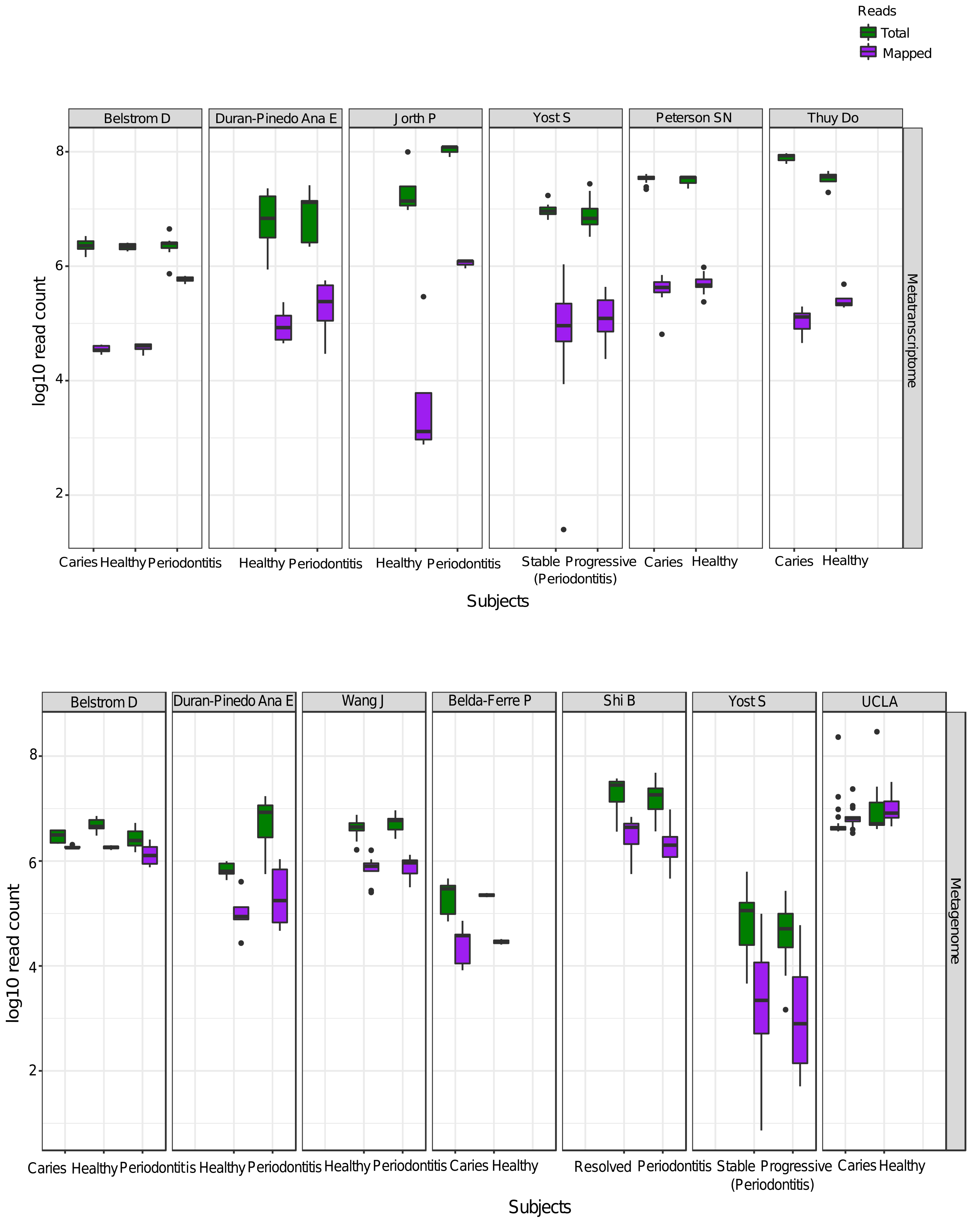

**Figure S10.**
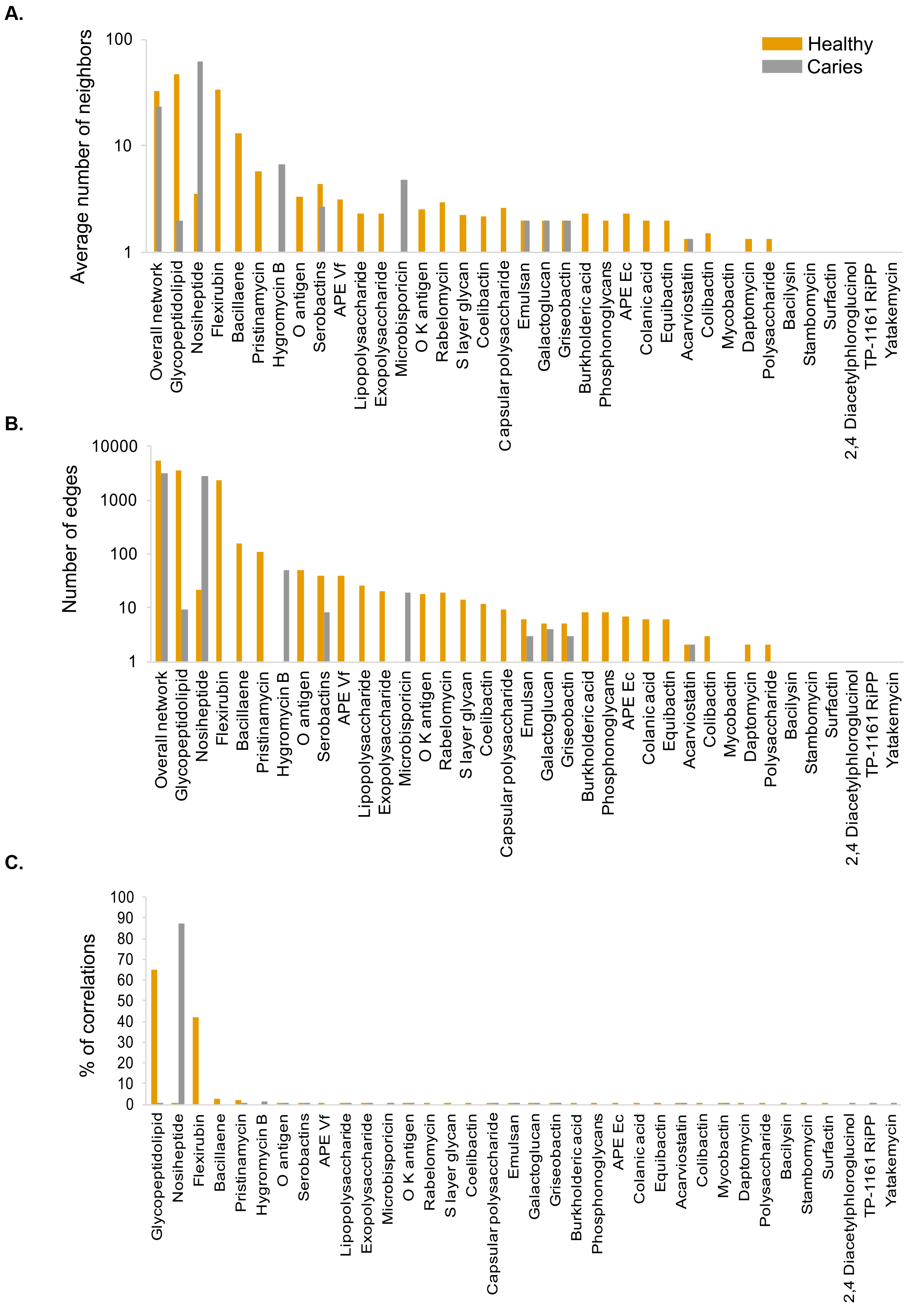

**Figure S11.**
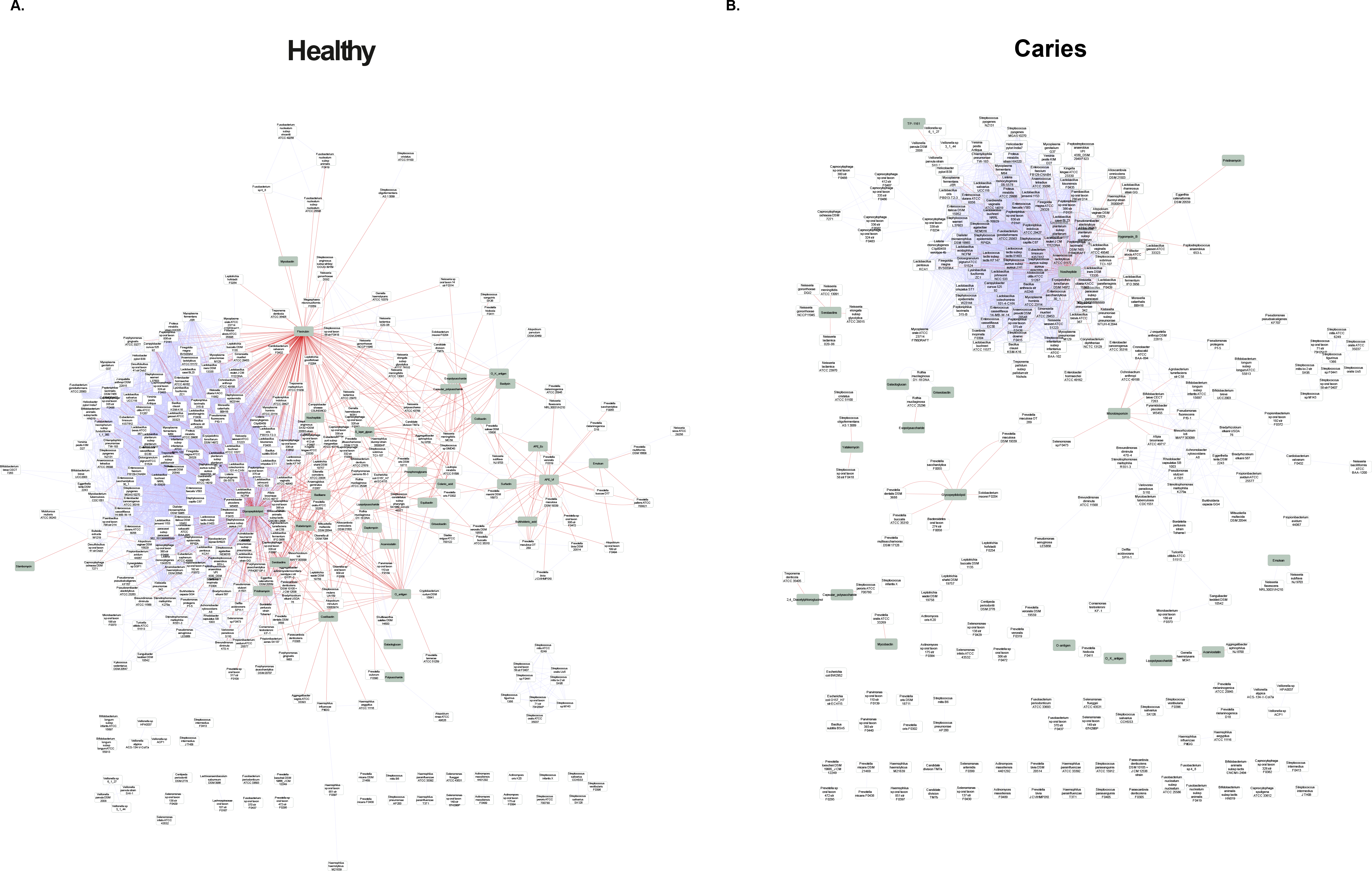

**Figure S12.**
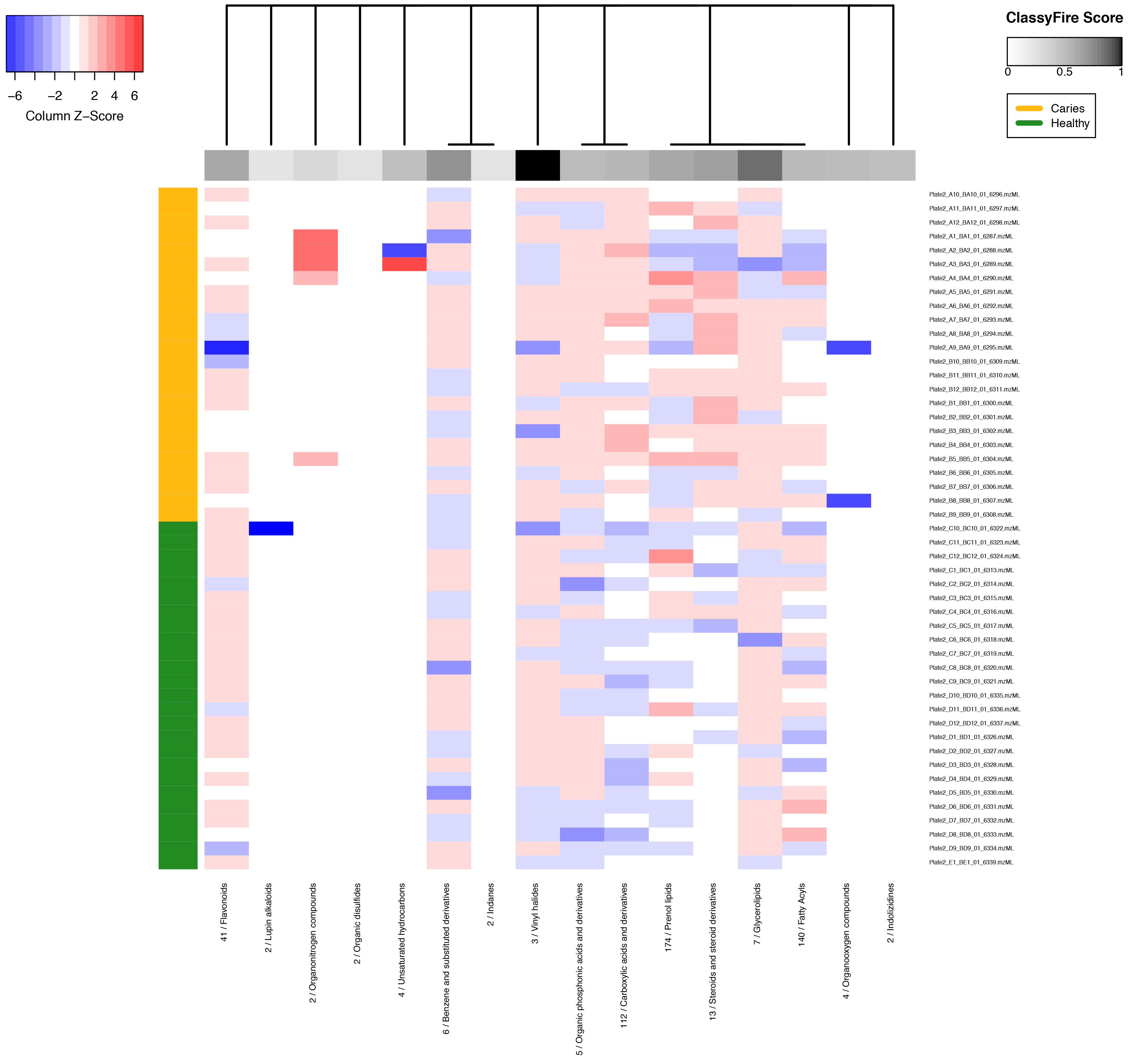

**Figure S13.**
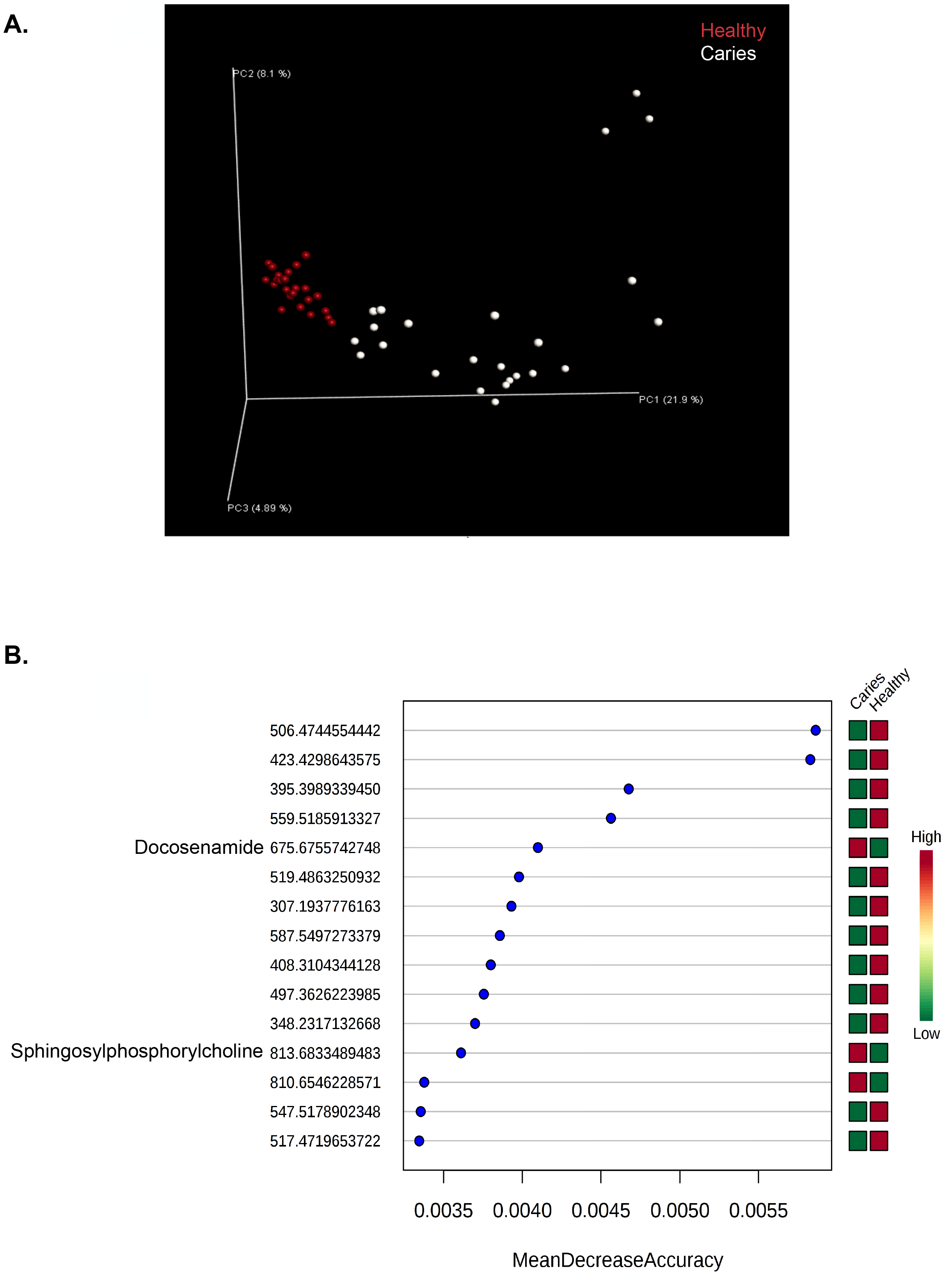

